# MicroRNA-122 supports robust innate immunity in hepatocytes by suppressing STAT3 phosphorylation

**DOI:** 10.1101/250746

**Authors:** Hui Xu, Shi-Jun Xu, Shu-Juan Xie, Yin Zhang, Jian-Hua Yang, Wei-Qi Zhang, Man-Ni Zheng, Hui Zhou, Liang-Hu Qu

## Abstract

The intrinsic innate immunity of hepatocytes is essential for the control of hepatitis viruses and influences the outcome of antiviral therapy. MicroRNA-122 (miR-122) is the most abundant microRNA in hepatocytes and is a central player in liver biology and disease. However, little is known about the role of miR-122 in hepatocyte innate immunity. Herein, we show that restoring miR-122 levels in hepatoma cells markedly increased the activation of both type III and type I interferons (IFNs) in response to hepatitis C virus (HCV) RNA or poly(I:C). We determined that miR-122 promotes IFN production through down-regulating the tyrosine (Tyr705) phosphorylation of STAT3. We show that STAT3 represses IFN activation by inhibiting interferon regulatory factor 1 (IRF1), which is rate-limiting for maximal IFN expression, especially type III IFNs. Through large-scale screening, we identified that miR-122 targets MERTK, FGFR1 and IGF1R, three oncogenic receptor tyrosine kinases that directly promote STAT3 phosphorylation. These findings reveal a previously unknown role for miR-122 in hepatic immunity and indicate a new potential strategy for treating hepatic infections through targeting STAT3.

## INTRODUCTION

Viral infection is a leading cause of liver diseases. The first line of immune defence against hepatitis viruses is the cell-intrinsic innate immunity within hepatocytes, which recognizes viruses as nonself and induces local antiviral defences in the infected cell and liver tissue that recruit and modulate the actions of the immune cells of the adaptive immune response^1^. It is now well-recognized that innate immunity not only plays a critical role in the initial containment of viraemia during acute infections but also continues to contribute throughout the course of a persistent viral infection^2^. Therefore, a greater understanding of the mechanism and regulation of innate immunity can pave the way for highly effective therapies.

Among the five hepatitis viruses, hepatitis C virus (HCV) affects more than 170 million people worldwide; most patients (80–85%) who become acutely infected cannot clear this virus, and the infection becomes chronic^3^. A substantial amount of data have shown that both spontaneous and therapy-induced HCV clearance are significantly associated with genetic variations in *IFNL3* and *IFNL4*, two interferon (IFN) genes of the type III IFN family (also known as IFN-λs) (summarized in reference 4)^4^. Very recently, the favourable *IFNL3* genotype has been suggested to increase the expression of IFNλ3 (also known as IL-28B)^4–6^. Since type III IFNs have a primary antiviral role^7, 8^ and constitute the dominant IFN subclass produced by hepatocytes in response to HCV infection^9–11^, these observations suggest that reinforcing innate IFN-signalling in infected cells may be an effective approach towards a cure for HCV.

MicroRNAs (miRNAs) are a large family of small non-coding RNAs that regulate various developmental and physiological processes^12^. While most miRNAs are ubiquitously expressed, a set of miRNAs are expressed with a higher degree of tissue specificity^13^ and are linked to specific functions in individual tissues or cell types^14^. MicroRNA-122 (miR-122) is a liver-specific miRNA that is exclusively expressed in hepatocytes^13^. Antisense-mediated inhibition^15, 16^ and knockout studies^17, 18^ demonstrate a critical role for this miRNA in the maintenance of liver homeostasis and hepatocyte function. Meanwhile, miR-122 is down-regulated or lost in human hepatocellular carcinoma (HCC)^19–21^, and delivering miR-122 mimics significantly inhibited the growth of HCC xenografts in nude mice^21, 22^, indicating that miR-122 serves as a tumour suppressor. Currently, the restoration of miR-122 levels has been suggested as a therapeutic approach in liver fibrosis and HCC development^23^. However, in contrast, miR-122 plays an undesirable role in the HCV life cycle^24^. miR-122 can be recruited to the 5′-end of HCV genomic RNA^24^, where it has a positive effect on viral translation, replication and stabilization^25–31^. Accordingly, antagonizing miR-122 as a novel approach for treating HCV infection has gained clinical interest^32, 33^.

Currently, although miR-122 has been recognized as a key player in liver biology and disease, whether miR-122 regulates hepatocyte innate immunity is unclear. Several important clues have implied the possible involvement of miR-122 in hepatic innate immunity. First, although miR-122 is required for the HCV life cycle, clinical observations have shown that deceased miR-122 levels in individuals with hepatitis C are associated with a poor response to interferon therapy^34, 35^, suggesting that miR-122 may promote the IFN response in infected hepatocytes. Second, the innate immune response is robust in primary hepatocytes but is strongly impaired in most hepatoma cell lines (in which miR-122 is significantly down-regulated or lost)^36, 37^. Third, miR-122 has an antiviral role in HBV infection^38, 39^; perhaps this effect is achieved partially by modulating the innate immunity. Finally, given that miR-122 is a formidable tumour suppressor and its loss results in hepatitis^17, 18^, this miRNA may take part in the regulation of hepatic immunity. Our current study was initially aimed at investigating if miR-122 regulates the hepatocyte IFN response to HCV RNA. We were surprised to find that miR-122 markedly enhanced the IFN response to HCV but was not specific to HCV. Importantly, we found that miR-122 promotes IFN expression by down-regulating STAT3 phosphorylation, which inhibits IRF1 and represses the transcriptional activation of IFNs. Therefore, our findings have identified a previously undiscovered network that links miR-122 to hepatic immunity through STAT3.

## RESULTS

### miR-122 enhances the hepatocyte IFN response to HCV RNA

To select a stable and reliable cell model for our study, we examined the functional innate immunity of three human hepatoma-derived cell lines: HepG2, Huh7 and Huh7.5.1^40^. Since HCV RNA is the predominant stimulator of the IFN response^41^, we transfected JFH1^42^ HCV genome RNA into these cell lines and then measured the induction of STAT1 phosphorylation (p-STAT1), a marker of IFN signalling activation. While HepG2 cells exhibited drastic p-STAT1 (Tyr701) induction compared to that of Huh7 cells, no p-STAT1 induction was observed in the Huh7.5.1 cells (Supplementary Fig. 1a). Analysis of melanoma differentiation-associated protein 5 (MDA5), an interferon-stimulated gene (ISG) that is markedly induced upon IFN signalling activation, gave similar results (Supplementary Fig. 1a). These results are consistent with previous reports showing that HepG2 cells have moderate innate immunity compared to that of primary hepatocytes^36^; Huh7 has minimal innate immunity due to a lack of Toll-like receptor 3 (TLR3)^36^; and Huh7.5.1 harbours additional notable defects in innate immunity due to a mutational inactivation of RIG-I^43^. Analysis of IFN mRNA levels by quantitative reverse transcription-polymerase chain reaction (qRT-PCR) revealed that the HCV RNA induced relatively robust IFN activation, specifically of type III IFNs (IL-29 and IL-28), in HepG2 cells (Supplementary Fig. 1b), which is similar to observations in primary human hepatocytes and *in vivo* studies^9–11^. Therefore, HepG2 may be a suitable cell model for studying innate hepatocyte immune responses.

To determine whether miR-122 is involved in hepatocyte immunity to HCV, we transfected miR-122 or control (miR-NC) mimics into HepG2 cells 48 hours (h) before JFH1 RNA transfection and then assessed the effect of miR-122 on IFN responses. miR-122 mimic transfection in HepG2 cells restored miR-122 expression to a level close to that of normal human liver (Fig. 1a). Consistent with the previous findings that miR-122 promotes HCV translation, an substantial increase in the expression of HCV core and non-structural 3 (NS3) proteins was observed in miR-122-transfected groups (Fig. 1b). Surprisingly, miR-122 originally inhibited p-STAT1 but dramatically enhanced p-STAT1 induction after HCV RNA stimulation, with the most significant difference at 24 h post HCV transfection (Fig. 1b). The MDA5 results were similar (Fig. 1b). The analysis of IFN mRNA levels revealed that miR-122 strikingly increased not only IFN-λs (IL-29 increased >300-fold, IL-28 increased >100-fold) but also IFN-β (increased >200-fold) (Fig. 1c). Analysis of IFN protein levels by ELISA revealed that IL-29/IL-28B levels were increased 9-fold and that the IFN-β level was undetected in the control group but increased to ~100 pg/ml in the miR-122-transfected group (Fig. 1d). We also performed the same assays in Huh7 cells. Although not highly significant, miR-122 increased p-STAT1 (Supplementary Fig. 1c) and IFN induction (Supplementary Fig. 1d) upon HCV RNA stimulation. In contrast, miR-122 did not promote the induction of p-STAT1 in HepG2 cells treated with either IFN-β or IL-29 (Fig. 1e), suggesting that miR-122 specifically enhances the transcriptional activation of IFNs in hepatocytes.

**Figure 1.**
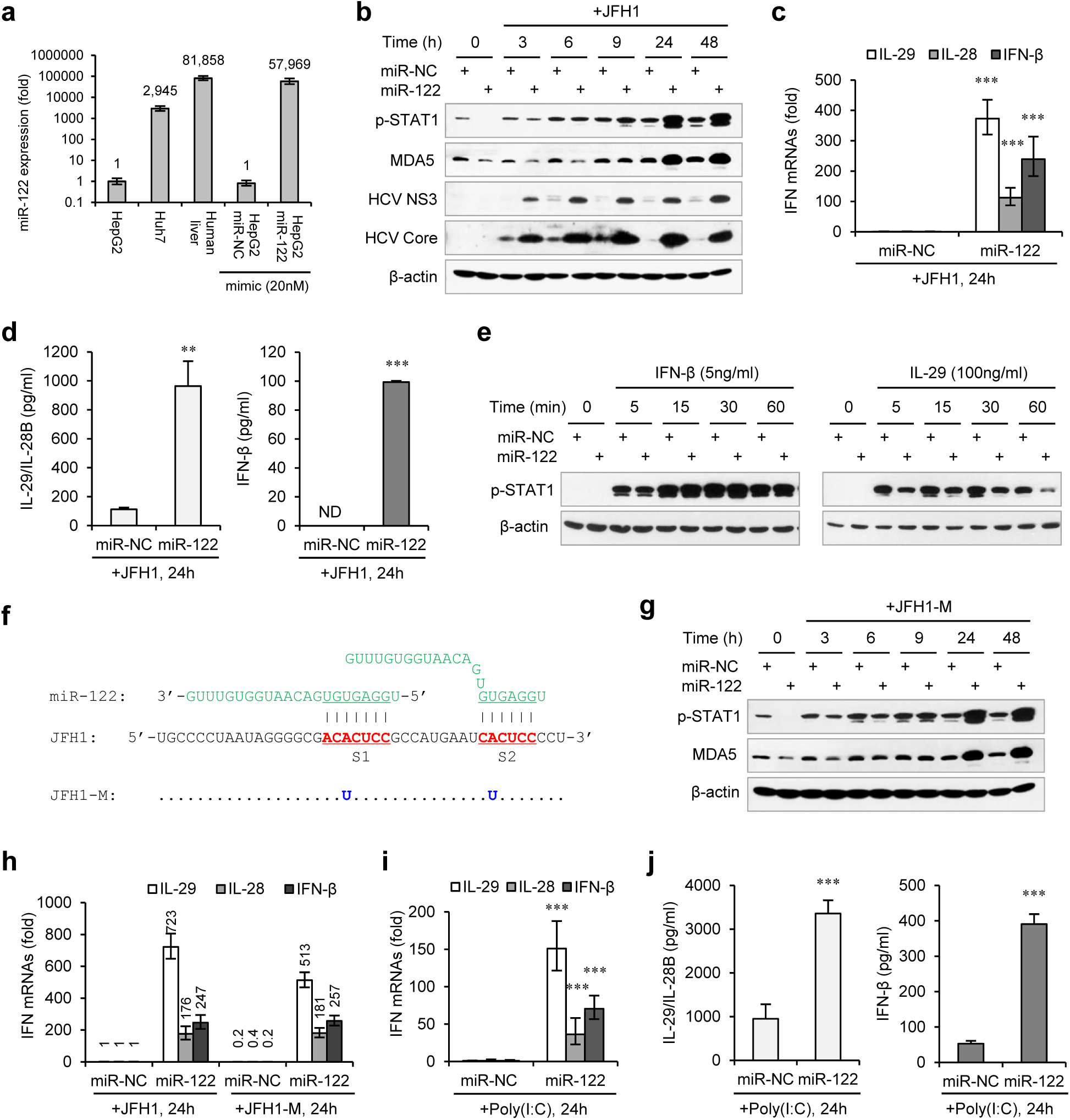
miR-122 enhances the activation of both type III and type I IFNs in HepG2 cellsstimulated with HCV RNAs or poly(I:C). (**a**) qRT-PCR analysis of miR-122 in hepatoma cell lines and in a normal liver tissue. The relative expression of miR-122 was normalized to U6 in each sample and then compared with the miR-122 level in HepG2. (**b**) Western blot analysis of p-STAT1 (Tyr701), MDA5, HCV Core and NS3 in HepG2 cells first treated with miR-122 or negative control (miR-NC) mimics for 2 days, and then transfected with JFH1 RNA for 3-48 hours. (**c**, **d**) Analysis of type III (IL-29, IL-28) and type I (IFN-β) IFN mRNA expression by qPCR (**c**) and protein production by ELISA (**d**) in HepG2 cells transfected as in and harvested at 24 hours post-transfection of JFH1 RNA. In qPCR, IL-28A and IL-28B were detected by the same pair of primers and thus referred as IL-28. In ELISA, IL-29 and IL-28B were detected by the same set of antibodies. The relative expression of IFN mRNAs was normalized to GAPDH and then compared with the levels in miR-NC groups. N.D., not detected. (**e**) Analysis of p-STAT1 expression in HepG2 cells first transfected with mimics (NC or miR-122) for 2 days and then treated with IFN-β or IL-29 for 5-60 minutes. (**f**) Alignment of miR-122 to the binding sites (S1 and S2) in the 5′ -end of JFH1 and mutant JFH1 (JFH1-M). (**g**) Analysis of p-STAT1 and MDA5 treated with JFH1-M RNA (as in **b**). (**h**) qRT-PCR analysis of IFN mRNAs treated with either JFH1 or JFH1-M, as in **c**. (**i**, **j**) Analysis of IFN mRNAs (**i**) and proteins (**j**) treated with poly(I:C). qRT-PCR and ELISA data are shown as the mean ±SEM. \**P* < 0.05, \*\**P* < 0.01 and \*\*\**P* < 0.001.

### miR-122 enhances the IFN response to mutant HCV or poly(I:C)

Considering that miR-122 can bind to the HCV 5′ UTR and enhance viral replication, the effect of miR-122 on IFN activation might be caused by an increase in HCV RNA abundance. However, the HCV RNA level was only slightly higher (~ 1.5-fold) in miR-122-treated cells than in control-treated cells (Supplementary Fig. 1e), suggesting that this assumption is incorrect. We then compared the IFN response to full-genomic (JFH1) and subgenomic HCV (SGR-JFH1)^42^ RNA levels and found that the effect of miR-122 on SGR-JFH1-induced IFNs (Supplementary Fig. 1f) was similar to the effect on JFH1-induced IFNs (Fig. 1c), suggesting that miR-122 regulates IFNs independent of infectious virus production. To determine whether the miR-122-induced promotion of IFN activation depends on miR-122 binding to the HCV 5′ UTR, we studied a viral RNA mutant (JFH1-M) with single base substitutions in S1 and S2 (Fig. 1f) that ablate miR-122 binding^30^. Remarkably, the mutant viral RNA also induced robust IFN synthesis and p-STAT1, which were further enhanced by miR-122 (Fig. 1g, h), indicating that binding to HCV RNA is not required for the effect of miR-122 on IFNs. Furthermore, miR-122 also enhanced the IFN response to poly(I:C), a synthetic analogue of double-stranded RNA (dsRNA) (Fig. 1i, j, Supplementary Fig. 1 g), indicating that the effect of miR-122 on the IFN response is not specific to HCV but is functional with other viruses.

### miR-122 promotes the IFN response by suppressing STAT3 phosphorylation

To understand the mechanism by which miR-122 regulates IFN activation, we performed reporter assays on four immune signalling pathways, including nuclear factor kappa B (NF-κB), interferon regulatory factor 1 (IRF1), type I IFN (using interferon-stimulated response element, ISRE), and STAT3. Consistent with the up-regulation of IFN production and p-STAT1 in the above results, miR-122 up-regulated ISRE promoter activity 24 h after SGR-JFH1 RNA treatment (Supplementary Fig. 2a). miR-122 also up-regulated the activity of IRF1, a key transcriptional regulator of the IFN response (Supplementary Fig. 2a). In contrast, miR-122 significantly repressed STAT3- and NF-κB-responsive promoters both before and after HCV RNA transfection (Fig. 2a, Supplementary Fig. 2a).

**Figure 2.**
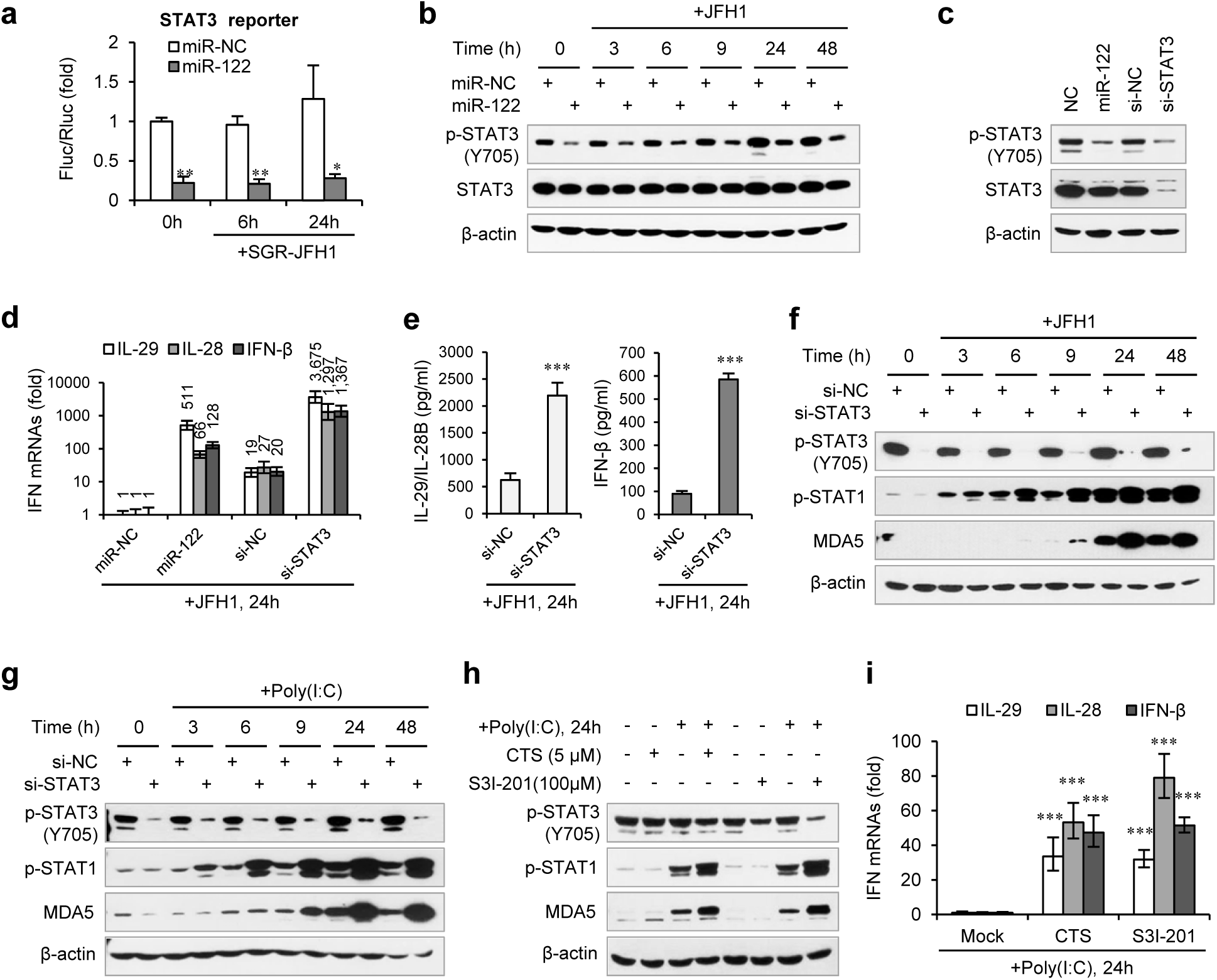
miR-122 enhances IFN response by suppressing p-STAT3. (**a**) Luciferase activity of STAT3 responsible promoter construct in HepG2 cells co-transfected with mimics (NC or miR-122) for 2 days, and then transfected with SGR-JFH1 RNA for the indicated time. The relative luciferase activities are the ratio of Firefly/Renilla luciferase normalized to that in cells transfected with NC mimic without SGR-JFH1 stimulation. (**b**) Western blot analysis of total and phosphorylated STAT3 (p-STAT3, Tyr705) in HepG2 cells first treated with mimics (NC or miR-122) for 2 days, and then transfected with JFH1 RNA. (**c**) Analysis of STAT3 protein in HepG2 cells treated with NC mimic, miR-122 mimic, NC siRNA or STAT3 siRNA (a mix of three independent siRNAs, 20nM in total). (**d**) qRT-PCR analysis of IFN mRNAs in HepG2 cells first treated with miR-122 mimics or STAT3 siRNAs for 2 days, and then transfected with JFH1 RNA. The relative expression of IFN mRNAs was normalized to GAPDH and then compared with the levels in miR-NC groups. (**e**, **f**) Analysis of IFN proteins (**e**) and p-STAT1 induction (**f**) in HepG2 cells transfected with STAT3 siRNA and then JFH1 RNA. (**g**) Analysis of p-STAT1 induction in HepG2 cells transfected with STAT3 siRNA and then poly(I:C). (**h**, **i**) Analysis of p-STAT1 protein and IFN mRNAs in HepG2 cells treated with either S3I-201 or Cryptotanshinone (CST) for 24 hours, and then transfected with poly(I:C). Luciferase data are shown as the mean + SD. qRT-PCR and ELISA data are shown as the mean ± SEM. \**P* < 0.05, \*\**P* < 0.01 and \*\*\**P* < 0.001.

While NF-κB activation is required for IFN induction^44^, STAT3 negatively regulates type I IFN-mediated antiviral responses^45, 46^. We first tested if miR-122 down-regulated the expression of STAT3. Intriguingly, miR-122 strongly inhibited STAT3 phosphorylation on a critical tyrosine residue (Tyr705) but did not affect the total STAT3 protein level, independent of HCV RNA stimulation (Fig. 2b). Although miR-122 appears to slightly down-regulate STAT3 mRNA expression, this alteration was extremely mild (Supplementary Fig. 2b). Comparing the effects of miR-122 overexpression and small interfering RNA (siRNA)-mediated STAT3 knockdown confirmed the specific regulation of miR-122 on phosphorylated STAT3 (Fig. 2c, Supplementary Fig. 2c). Next, we analysed the IFN response following STAT3 knockdown and HCV RNA stimulation. Excitingly, STAT3 knockdown greatly increased IFN production and p-STAT1 induction (Fig. 2d, e, f). Notably, the IFN level was significantly higher in the STAT3 knockdown cells than in the miR-122-treated cells (Fig. 2d). The analysis of the effects of STAT3 knockdown on p-STAT1 or IFN activation using JFH1-M or poly(I:C) revealed similar results (Fig. 2 g, Supplementary Fig. 2d, e). Furthermore, blocking STAT3 phosphorylation by the chemical inhibitor S3I-201^47^ or the natural compound cryptotanshinone (CTS)^48^ (Supplementary Fig. 2f) also significantly increased the IFN response (Fig. 2h, i). Therefore, these data demonstrate that miR-122 regulates IFN activation via repressing STAT3 phosphorylation.

### STAT3 inhibits IRF1 by binding to IRF1 promoters and enhancers

To understand how STAT3 regulates IFN transcriptional activation, we analysed the expression of five transcription factors responsible for IFN activation: IRF1, IRF3, IRF7, RELA and NFKB1 (also known as NF-κB p65 and p50). Both miR-122 overexpression and STAT3 knockdown substantially increased IRF1 expression at 24 h post HCV RNA transfection but did not significantly impact the other expression levels (data on IRF7 is not shown because it could be hardly detected) (Fig. 3a). Analysis of the protein expression of IRF1 and IRF3 at different times after HCV RNA stimulation showed that STAT3 knockdown increased IRF1 activation very early or even in unstimulated cells (Fig. 3b), suggesting that IRF1 up-regulation might account for the increased IFN activation. As expected, IRF1 overexpression directly triggered IFN expression (Fig. 3c) and STAT1 phosphorylation (Fig. 3d). Notably, IRF1 overexpression mainly induced IFN-λ expression (Fig. 3d), which is consistent with a recent report showing that IRF1 is specifically required for IFN-λ activation^49^. Moreover, IRF1 induced STAT1 phosphorylation and MDA5 expression in a dose-dependent manner, and even a small amount of IRF1 significantly increased p-STAT1 and MDA5 expression (Fig. 3e). Comparing IRF1 activity to that of six known transcriptional regulators of IFN-β^50^ also revealed that IRF1 is the major inducer of p-STAT1 (Fig. 3f). A previous study suggested that STAT3 sequesters STAT1 into STAT1:STAT3 heterodimers and thus inhibits STAT1-dependent IRF1 activation^45^. However, in our system, STAT3 knockdown did not increase the IRF1 induction by either IFN-β or IL-29 (Fig. 3g), suggesting that STAT3 mainly represses HCV-induced IRF1 activation.

**Figure 3.**
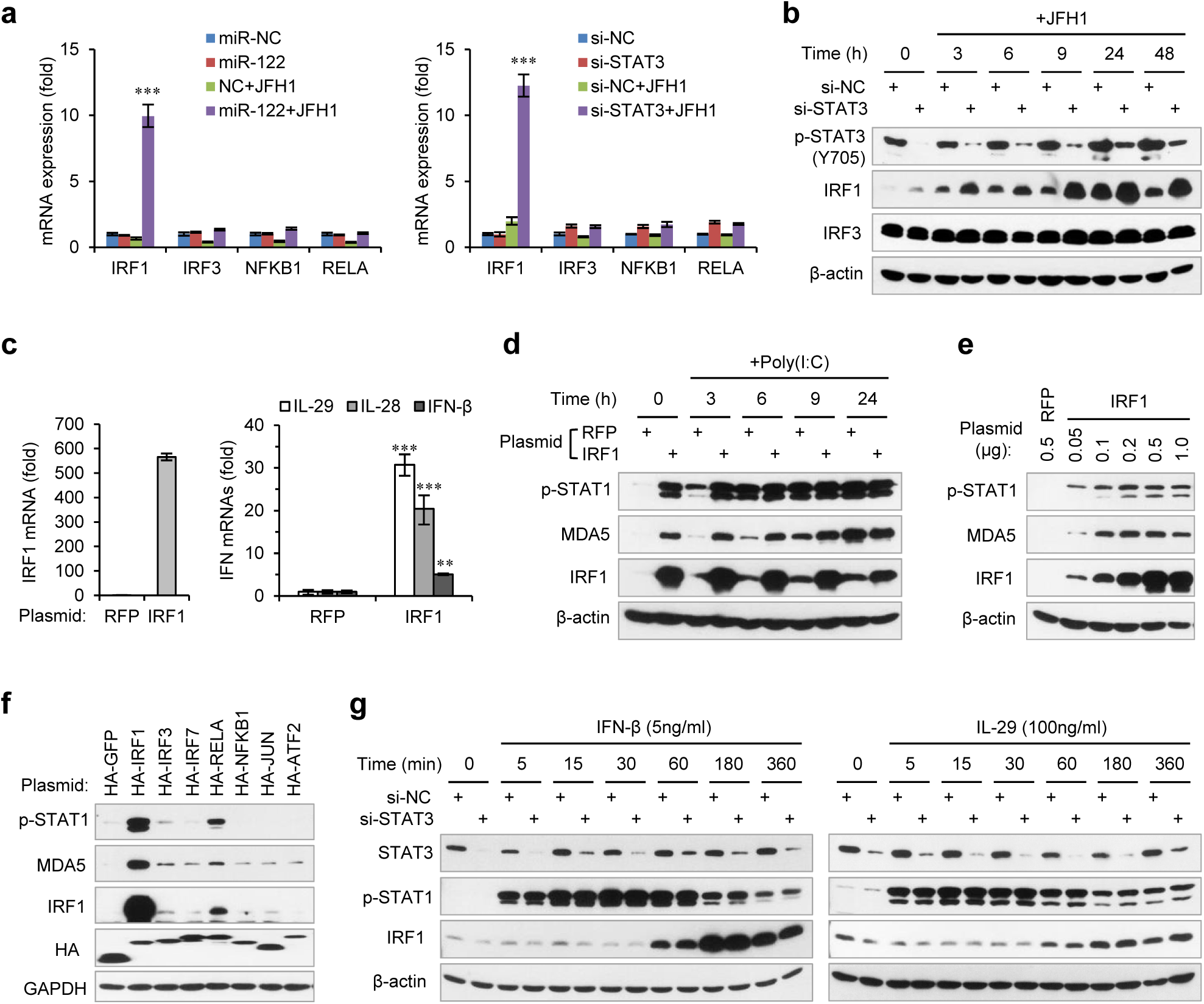
STAT3 inhibits HCV RNA-induced transcriptional activation of IRF1. **(a)** qRT-PCR analysis of IRF1, IRF3, NFKB1 and RELA in HepG2 cells first treated with mimics (left) or siRNAs (right), and then transfected with or without JFH1 RNA for 24 hours. The relative expression of mRNAs was normalized to GAPDH and then compared with the levels in miR-NC or si-NC samples without JFH1 RNA treatment. (**b**) Analysis of IRF1 and IRF3 protein expression in HepG2 cells transfected with STAT3 siRNA and then JFH1 RNA. (**c**) qRT-PCR analysis of IRF1 and IFNs in HepG2 cells transfected with IRF1 plasmid for 2 days. A plasmid expressing RFP was used as negative control. (**d**) Analysis of p-STAT1 and MDA5 in HepG2 cells transfected with IRF1 plasmid for 2 days, and then treated with poly(I:C) for 3-24 hours. (**e**) Analysis of p-STAT1 and MDA5 in HepG2 cells transfected with indicated dose of IRF1 plasmids (0.05-1 μg/well for 24-well-plate) for 2 days. (**f**) Analysis of IRF1, p-STAT1 and MDA5 in HepG2 cells transfected with plasmids expressing 7 HA-tagged transcription factors (for 2 days). HA-GFP was used as a negative control. (**g**) Analysis of IRF1 and p-STAT1 in HepG2 cells first transfected with STAT3 siRNA for 2 days, and then treated with IFN-β or IL-29 for 5-360 minutes. qRT-PCR data are shown as the mean ± SEM. \**P* < 0.05, \*\**P* < 0.01 and \*\*\**P* < 0.001.

Genome-wide chromatin immunoprecipitation sequencing (ChIP-seq) data from the ENCODE project have revealed seven STAT3 binding clusters (BS1 to BS7) on the IRF1 gene locus, and there are three conserved binding motifs for STAT3 within BS1, BS3 and BS4 (Fig. 4a, Supplementary Fig. 3a, b, Supplementary Table 1). Because these ChIP-seq data were obtained in cell lines other than hepatocytes, we performed ChIP experiments to determine if these regions were bound by STAT3 in HepG2 cells. As expected, STAT3 appeared to bind to four sites (BS1, BS2, BS3 and BS4) and weakly bound to the other three sites (Fig. 4b). Interestingly, using an antibody that specifically recognizes p-STAT3(Tyr705), we found that p-STAT3 bound to BS2, BS3 and BS4 but not BS1 (Fig. 4b). By comparison, RELA, an activator of IRF1 (Fig. 3f), only weakly bound to BS1, BS2 and BS4 (Fig. 4b). These data indicate that STAT3 can directly bind several sites on the IRF1 gene.

**Figure 4.**
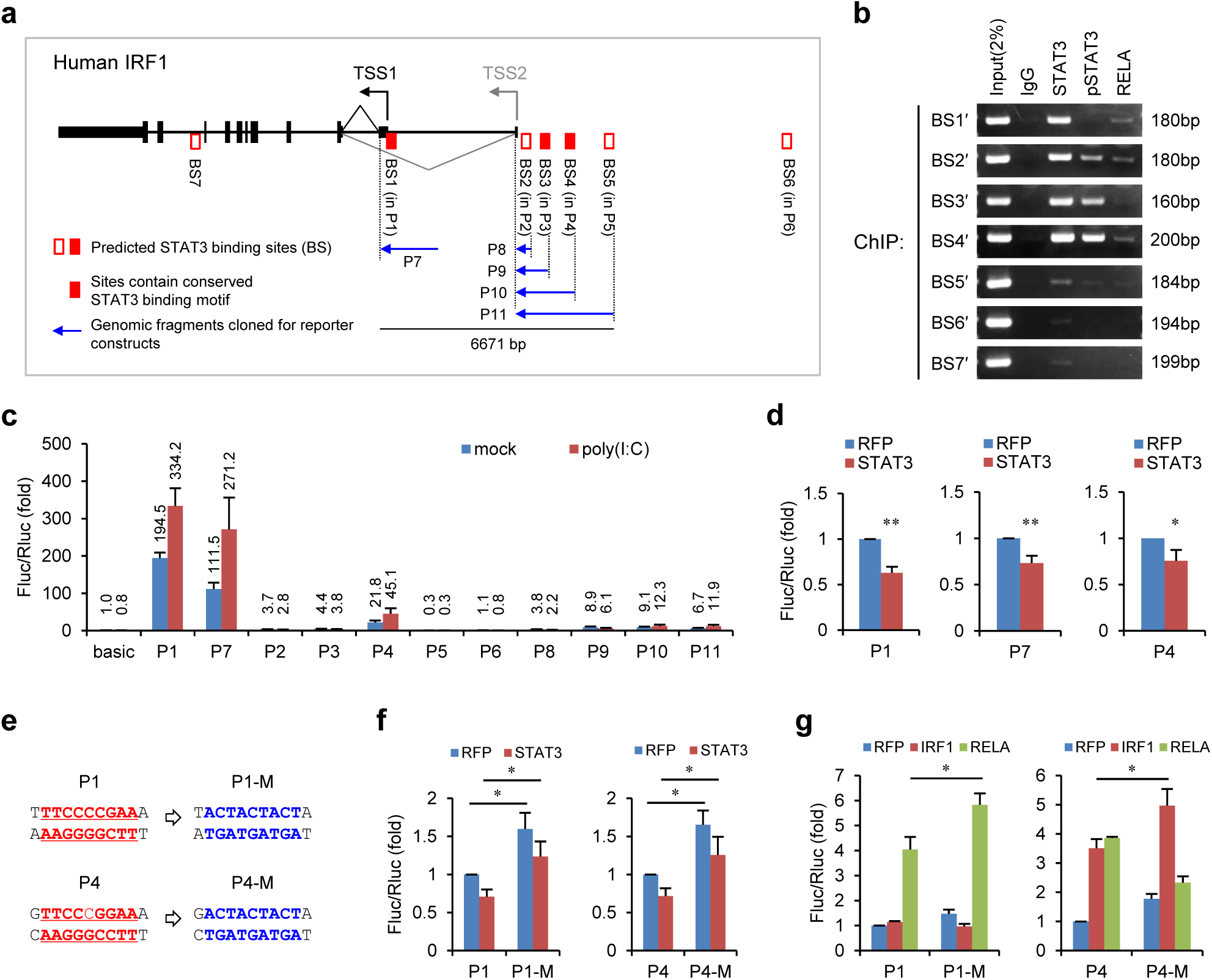
STAT3 binds to the promoter and enhancers of IRF1 gene. (**a**) Schematic representation of STAT3 binding sites (BS1-BS7, STAT3 ChIP-seq clusters identified by the ENCODE project) on human IRF1 gene. The DNA fragments selected for reporter constructs (P1-P11) are also shown. (**b**) ChIP-PCR assays show the binding of STAT3, p-STAT3 and RELA on the selected gene fragments of IRF1. BS1′-BS7′ are short fragment (160-200 bp) corresponding to BS1-BS7 ChIP-seq clusters (290-410 bp), respectively. (**c**) Luciferase activity of different IRF1 promoter/enhancer constructs (P1 to P11) in HepG2 cells treated with or without poly(I:C) (for 24 hours). The relative luciferase activities are the ratio of Firefly/Renilla luciferase normalized to the control construct (basic) without poly(I:C) stimulation. (**d**) Luciferase activity of P1, P7 and P4 constructs in 293FT cells co-transfected with STAT3 or control (RFP) plasmids. (**e**) The sequences of the STAT3 binding motifs in wild-type (P1, P4) constructs and mutant (P1-M, P4-M) constructs. (**f**, **g**) Luciferase activity of IRF1 constructs in 293FT cells co-transfected with STAT3 plasmid (**f**) or IRF1/RELA plasmids (**g**). The relative luciferase activities were normalized to that in cells co-transfected with RFP plasmid. Luciferase data are shown as the mean ± SE. \**P* < 0.05 and \*\**P* < 0.01.

To determine which sites are critical for IRF1 regulation, we generated a series of luciferase reporter constructs (Fig. 4a) and assessed their activities. Constructs P1 to P6 each harbour a DNA fragment that completely includes the BS1 to BS6 clusters, respectively. Constructs P7 to P11 contain longer DNA inserts that cover one or more ChIP-seq clusters. Consistent with previous studies demonstrating that the IRF1 promoter is located immediately upstream of the IRF1 transcriptional start site (TSS1 in Fig. 4a)^51^, the activities of the P1 and P7 constructs (both constructs contain the putative promoter sequences) were substantially higher than those of all the other constructs (Fig. 4c). Although there is another TSS (TSS2) for IRF1, the activities of constructs P8 to P11 were relatively low in HepG2 cells (Fig. 4c), suggesting that this alternative TSS may be minimally utilized in HepG2 cells. Notably, the activities of constructs P1, P7, P4, P10 and P11 were significantly increased by poly(I:C) transfection, whereas the others were not (Fig. 4c), indicating that activating elements are located within P1 (BS1) and P4 (BS4).

To assess the effect of STAT3 on P1, P7 and P4, constructs were co-transfected into 293FT cells with a STAT3-expressing plasmid. As expected, STAT3 inhibited the activities of all three constructs (Fig. 4d). Next, we performed mutational analyses on the predicted STAT3 binding sites within P1 and P4 (Fig. 4e). Although the STAT3 binding site mutations did not completely abolish the repressing effect of STAT3 on P1 or P4, the activities of the mutant constructs (P1-M, P4-M) were significantly increased (Fig. 4f), suggesting that STAT3 may repress the activities of P1 and P4 partially through binding to these two sites. In addition, we also analysed if the mutations in the STAT3 binding sites would increase the activation of the P1 and P4 constructs. Consistent with the observation that RELA overexpression significantly promotes IRF1 expression (Fig. 3f), there is a conserved NF-κB binding site in P1 (Supplementary Fig. 4a). In addition, there is a conserved IRF binding site in P4 (Supplementary Fig. 4b), suggesting that IRF1 may also regulate its own transcription. As expected, mutating the STAT3 binding sites significantly increased the activating effects of RELA and IRF1 on P1 and P4, respectively (Fig. 4g). Interestingly, STAT3 binding site mutations in P4-M blocked the activating effects of RELA on P4, probably due to the mutation also influencing RELA binding to non-canonical NF-κB binding sites (Supplementary Fig. 4b). Taken together, these results demonstrate that STAT3 represses IRF1 activation by binding to the promoter and enhancers of the IRF1 gene.

### A group of genes mediated miR-122 regulation of STAT3 and IFNs

We first examined whether miR-122 inhibited well-characterized activators of STAT3, including IL-6, IL-6R, gp130 (encoded by *IL6ST*) and EGFR^52^. Although miR-122 inhibited gp130 protein expression, miR-122 increased the expression of IL6R and EGFR, especially that of IL-6 (Supplementary Fig. 5a, b). Knocking down gp130 only slightly increased IFN activation (Supplementary Fig. 5c). In addition, we found that gp130 is downstream of STAT3 (Supplementary Fig. 5d). Thus, miR-122 may regulate some undefined regulators of p-STAT3.

To search for the genes that mediate miR-122 regulation of p-STAT3, we first performed microarrays to obtain the genes repressed by miR-122 and then screened for STAT3 activators (Fig. 5a). To obtain the genes truly regulated by miR-122, we used both transient and stable miR-122 overexpression in HepG2 cells. The transient mimic transfection was performed as above. For stable miR-122 overexpression, we generated a miR-122-Tet-On cell line in which high miR-122 expression level (~10-fold of that in Huh7 cells) can be induced by doxycycline treatment (Supplementary Fig. 6 a, b, c). In total, expression signals from 1349 probe sets (corresponding to 1141 genes) were down-regulated more than 20% by miR-122 in both groups (Fig. 5a, Supplementary Table 2). According to several references, we collected a total of 330 known and potential STAT3 activators that were detected in our microarray (Supplementary Table 3). Among the 330 candidates, 25 genes were down-regulated by miR-122 (Supplementary Table 4). The qRT-PCR data confirmed that 20 genes were significantly down-regulated by miR-122 (Fig. 5b), but the remaining five genes were hardly detected or were unchanged (data not shown).

**Figure 5.**
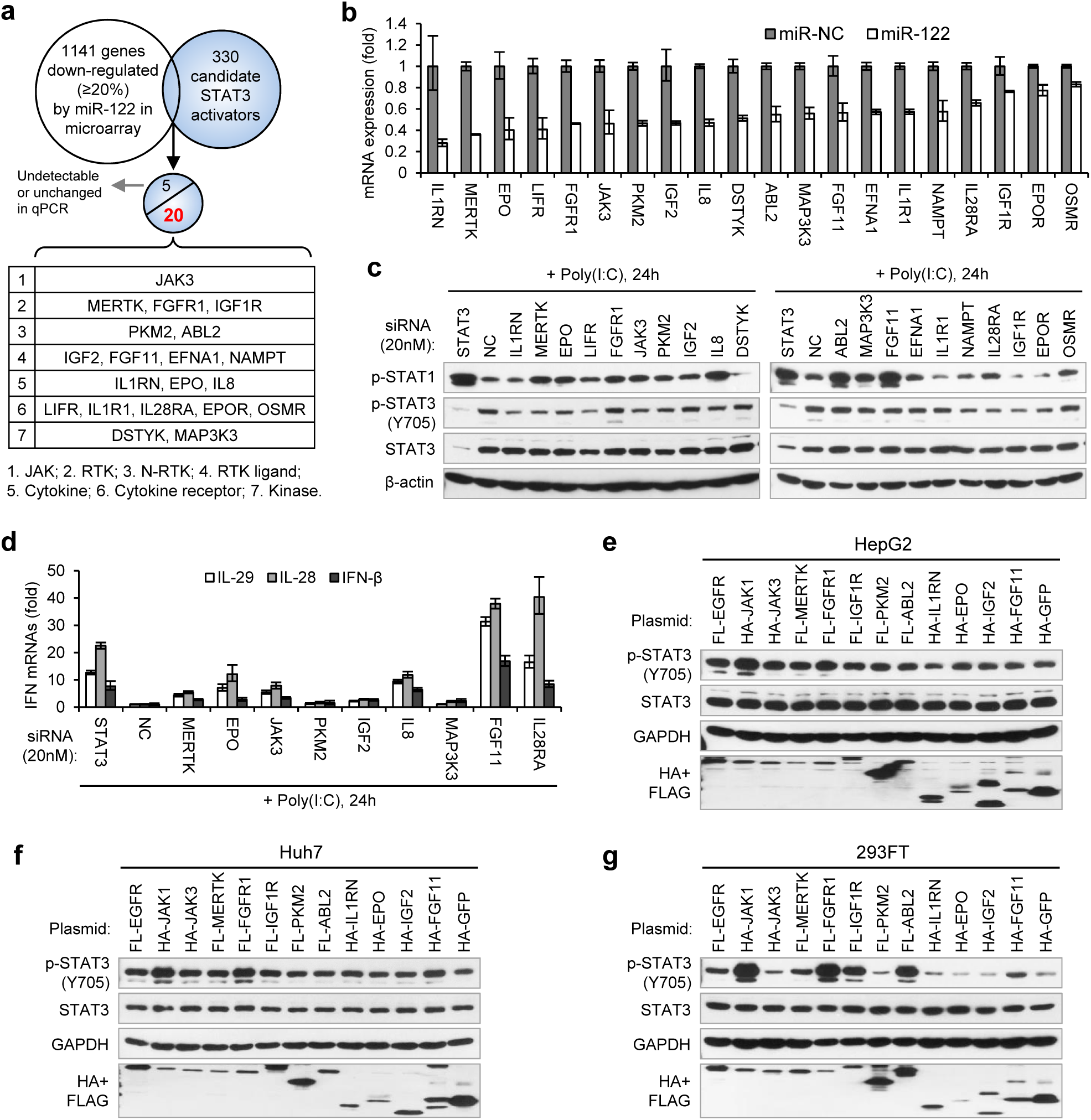
Identification of genes mediated miR-122 regulation of STAT3 and IFNs. (**a**)Strategy for identifying genes that mediated miR-122 regulation of STAT3. (**b**) qRT-PCR analysis of the 20 genes in HepG2 cells transfected with miR-122 or NC mimics (50 nM). The expression of each gene was normalized to its level in miR-NC-treated cells. The genes are ranked by the repression ratio. (**c**) Analysis of p-STAT1, p-STAT3 and total STAT3 protein in HepG2 cells first treated with indicated siRNAs (20 nM) for 2 days, and then transfected with poly(I:C) for 24 hours. (**d**) qRT-PCR analysis of IFNs in HepG2 cells treated with siRNAs and poly(I:C), as in **c**. (**e-g**) Analysis of total and phosphorylated STAT3 in HepG2 (**e**), Huh7 (**f**) and 293FT (**g**) cells transfected with plasmids expressing HA- or Flag (FL)-tagged proteins. HA-JAK1 and FL-EGFR plasmids were employed as positive controls. HA-GFP was used as a negative control. qRT-PCR data shown are mean ± SEM.

To determine whether these genes mediated the effect of miR-122 on p-STAT3 and the IFN response, we first performed knockdown experiments (Supplementary Fig. 7). While knocking down most genes (except for FGFR1, DSTYK, ABL2 and OSMR) reduced the p-STAT3 level, the knockdown of ten genes (MERTK, EPO, FGFR1, JAK3, PKM2, IGF2, IL8, ABL2, MAK3K3 and FGF11) appeared to increase the p-STAT1 level (Fig. 5c). Notably, some of these genes (MERTK, EPO, JAK3, PKM2 and IL8) significantly affected both p-STAT1 and p-STAT3. The qRT-PCR data confirmed that p-STAT1 up-regulation was accompanied by an increase in IFN expression (Fig. 5d). Next, we performed overexpression experiments on ten genes, which were selected for because their knockdown either significantly increased p-STAT1 levels (FGFR1, IGF2, ABL2 and FGF11), significantly reduced p-STAT3 levels (IL1RN and IGF1R), or both (MERTK, EPO, JAK3 and PKM2). JAK1 and EGFR, two well-characterized STAT3 activators, were employed as positive controls. Most likely, because STAT3 was constitutively phosphorylated in HepG2 cells, the overexpression of most of these genes did not further increase the p-STAT3 level in HepG2 cells (Fig. 5e). However, in Huh7 or 293FT cells, the overexpression of most of the genes increased p-STAT3 levels (Fig. 5f, g). In particular, FGFR1, IGF1R, MERTK, JAK3 and FGF11 clearly up-regulated p-STAT3 levels, but the effect was slightly different in the two cell lines (Fig. 5f, g). Taken together, these results demonstrate that miR-122 may regulate STAT3 phosphorylation and the IFN response through repressing several STAT3 activators.

### miR-122 targets receptor tyrosine kinase (RTK) signalling

To further understand how miR-122 regulates p-STAT3 levels, we assessed the expression of all 20 genes in HepG2 and Huh7 cells and in a normal human liver sample. Interestingly, the mRNA expression of six genes (IGF1R, PKM2, FGFR1, FGF11, IGF2 and MERTK) was significantly higher (>2-fold) in the HepG2 cells than in the normal liver cells (Fig. 6a), suggesting that these genes might account for the persistent STAT3 activation in HepG2 cells. To determine which gene was most influential, we again performed knockdown experiments with a higher dose of siRNAs (50 nM). Remarkably, while PKM2 and IGF2 knockdown moderately decreased p-STAT3 levels, MERTK and IGF1R knockdown markedly decreased the p-STAT3 level (Fig. 6b), indicating that MERTK and IGF1R might be the key factors that mediate miR-122’s regulation on p-STAT3. Although FGFR1 and FGF11 knockdown alone did not significantly reduce the p-STAT3 level, in view of the overexpression experiment results (Fig. 5f, g), these genes might also be responsible for the constitutive phosphorylation of STAT3 in HepG2 cells.

**Figure 6.**
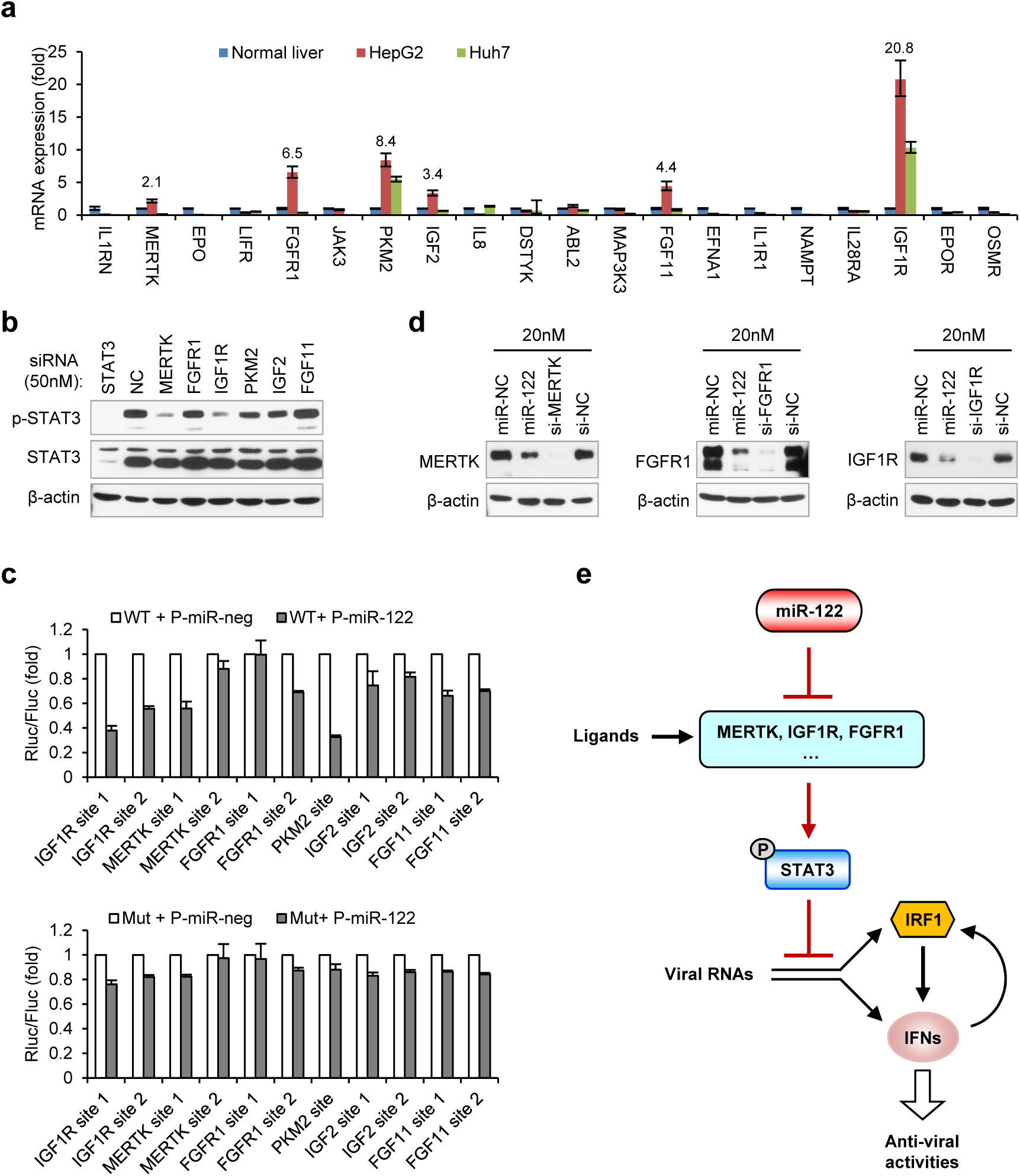
miR-122 targets RTK signalling to regulate STAT3 phosphorylation. (**a**) qRT-PCR analysis of the 20 genes in normal human liver, HepG2 and Huh7. The expression of each gene was normalized to its level in normal liver. (**b**) Analysis of total and phosphorylated STAT3 in HepG2 cells transfected with indicated siRNAs (50 nM) for 2 days. (**c**) Luciferase activity of wild-type (upper) and mutant (lower) target constructs in 293FT cells co-transfected with pcDNA6.2-miR-122 (P-miR-122) or pcDNA6.2-miR-neg (P-miR-neg) plasmids. The relative luciferase activities are the ratio of Renilla/Firefly luciferase normalized to that in P-miR-neg groups. (**d**) Analysis of MERTK, FGFR1 and IGF1R protein expression in HepG2 cells treated with miR-122 mimics or specific siRNAs (for 2 days). (**e**) Illustration of the mechanism by which miR-122 regulates STAT3 phosphorylation and IFN response. Phosphorylated STAT3 can inhibit IRF1 transcriptional activation and thus represses the induction of IFNs upon viral infection. In normal hepatocytes, by targeting three RTKs and other STAT3 activators, miR-122 strongly limits the phosphorylation of STAT3, enabling a robust IFN response upon infection.

Among these six genes, IGF1R^22^ and PKM2^53^ are known targets of miR-122. Using Targetscan^54^ or RegRNA^55^, we found that the remaining four genes also possess potential miR-122 binding sites (Supplementary Fig. 8). To investigate if miR-122 directly regulates these genes, we generated reporter constructs as previously described^56^ and performed luciferase assays. As positive controls, the miR-122 binding sites on IGF1R and PKM2 were also tested. Except for IGF2, all the other genes have at least one functional site that can be significantly repressed by miR-122 (Fig. 6c). In addition, mutating these sites largely or completely abolished the response to miR-122 (Fig. 6c), indicating the specificity of the repression. Consistent with the decreased mRNA expression, miR-122 overexpression significantly inhibited the protein expression of three RTKs: MERTK, FGFR1 and IGF1R (Fig. 6d). In conclusion, our results suggest that miR-122 suppresses STAT3 phosphorylation mainly through targeting RTK signalling.

## DISCUSSION

This study demonstrates that miR-122 promotes IFN-based innate immunity by regulating genes that contribute to the STAT3 phosphorylation level and thereby removes the negative regulation of STAT3 on IFN signalling. Our findings thus reveal a critical role for miR-122 in hepatocyte innate immunity, as illustrated in Fig. 6e. According to this model, miR-122 is responsible for restricting STAT3 phosphorylation to a low level in normal hepatocytes, which enables a robust innate immune defence upon infection.

It has been speculated that miR-122 is linked to each important aspect of hepatic function, as it is exclusively expressed in hepatocytes and constitutes 70% of the entire miRNA population in mature hepatocytes. The majority of the roles miR-122 plays in diverse hepatic functions and processes, such as cholesterol and lipid metabolism, hepatocyte differentiation and maturation, and liver tumour suppression, are easy to appreciate^23^. In contrast, the roles of miR-122 in hepatic infection and immunity are not very clear, as miR-122 appears to play almost completely different roles in HCV and HBV infections^23^. Our study provides the first direct evidence that miR-122 plays a general antiviral function in hepatocytes by enhancing the innate IFN signalling, which reveals an unknown aspect of the interplay between the virus, host and innate immune system. However, this role of miR-122 has gone undiscovered for a long time, probably because the role of miR-122 in innate immunity was largely masked by its unique requirement for HCV replication or because HCV partially sequestered the function of miR-122^57^. In addition, because HBV infection did not induce an obvious innate immune response, a role for miR-122 in innate immunity has not been observed in HBV infection, despite the fact that miR-122 inhibits HBV infections^38, 39^.

miR-122 plays an essential role in maintaining liver homeostasis, and the loss of miR-122 can lead to hepatosteatosis, hepatitis, and the development of tumours resembling HCC^17, 18^; however, this mechanism is not fully understood. Previous studies have shown that STAT3 activity is associated with the proliferation capacity of hepatocytes^58^, and STAT3 reactivation in primary human hepatocytes restores the ability to proliferate^59^. In addition, STAT3 is persistently activated in many cancers and plays a crucial role in the tumour initiation and malignant progression of many cancers by promoting pro-oncogenic inflammatory pathways and opposing anti-tumour immune responses^52^. Since miR-122 is frequently down-regulated or lost in HCC^19–21^, our findings suggest that the loss of miR-122 may be a critical factor causing constitutive STAT3 activation, which mediates dedifferentiation, inflammation and tumour immune-escape during cancer development.

Although a report suggested that miR-122 may directly target STAT3^60^, this potential direct regulation was not observed in our study. Instead, our results suggest that miR-122 represses STAT3 phosphorylation through targeting STAT3 activators, such as MERTK, FGFR1 and IGF1R. Notably, the overexpression of these RTKs also increased STAT1 phosphorylation (Supplementary Fig. 9). These results may explain why STAT1 is phosphorylated in HepG2 cells without HCV RNA treatment and why miR-122 initially inhibited p-STAT1 (Fig. 1b). Moreover, these results further suggest that miR-122 strongly limits the action of growth signals in normal hepatocytes by controlling the receptor levels, whereas in cells lacking miR-122, the growth signals are unlimited, resulting in the activation of both proliferation-promoting (downstream of RTKs) and inflammatory genes (downstream of the STAT proteins).

STAT3 is known to block the innate and adaptive immune responses in diverse cell types^45, 46, 61^, but this mechanism has remained elusive. Our results suggest that STAT3 strongly limits the IFN response, at least by repressing IRF1, a key transcriptional activator of IFNs. Intriguingly, when we transfected poly(I:C) into PC3 cells, a prostate cancer cell line that lacks STAT3 in the genome^62^, we found that the IFN activation was faster and stronger than that in HepG2 cells (Supplementary Fig. 10a). Moreover, plasmid DNA transfection into PC3 cells also resulted in high IFN production (Supplementary Fig. 10b). In summary, we hypothesize that STAT3 may function as an “antiknock reagent” in innate immunity by preventing an excessive IFN response. Important evidence supporting this view is the observation that STAT3 loss-of-function mutations result in Hyper-IgE Syndrome (or Job’s syndrome) in humans, an immunodeficiency syndrome involving an increased innate immune response^63, 64^.

Previous studies have revealed that STAT3 can be activated by different mechanisms during HCV infection^65, 66^. Large-scale screenings have also identified STAT3 as one of the main host factors required for HCV infection^67^. However, the key contribution of STAT3 to the HCV life cycle remains undefined. Our results suggest that the up-regulation of STAT3 activity may reduce the IFN response and thereby facilitate HCV infection. Interestingly, HCV can sequester miR-122 through a “sponge” effect and globally reduces miR-122 targets^57^, suggesting that HCV-induced miR-122 sequestration may be another mechanism that results in STAT3 activation and immune tolerance.

Our findings suggest miR-122 plays dual role in HCV infection. On the one hand, HCV replication depends on miR-122; and on the other hand, miR-122 promotes the IFN response to limit HCV infection. Consistent with the positive role of miR-122 in defence against HCV, deceased miR-122 levels in individuals with hepatitis C are associated with a poor response to IFN therapy^34, 35^. Liver biopsy studies on HCV-infected patients indicate that miR-122 expression is decreased significantly with fibrosis severity^68, 69^, suggesting that non-responders have a more serious infection than responders. Meanwhile, these observations suggest that HCV-induced chronic inflammation contributes to decreased miR-122 levels, which further blunts IFN signalling in HCV-infected hepatocytes. Therefore, the use of the anti-miRNA-122 oligonucleotide miravirsen to treat HCV in patients with serious infections should be carefully monitored in future clinical studies, as it may further impair the endogenous immune defence. Alternatively, our findings highlight the potential of STAT3 targeting for the treatment of HCV and other hepatitis viruses. In particular, targeting STAT3 will greatly reduce the risk of tumourigenesis and is suitable for patients with hepatic tumours.

## MATERIALS AND METHODS

### Mimics, siRNAs, antibodies, Reagents and Kits

miR-122 (4464066) and control (4464058) mimics were obtained from Ambion. Negative control siRNA were obtained from Dharmacon. Gene-specific siRNAs were obtained from RiboBio with three independent siRNAs for each gene. STAT3 siRNAs were: si-1 (GCCUCUCUGCAGAAUUCAA), si-2 (AGUCAGGUUGCUGGUCAAA) and si-3 (CCGUGGAACCAUACACAAA). Antibodies used are: Phospho-STAT1 (Tyr701, CST, 7649), Phospho-STAT3 (Tyr705, CST, 9145, for western blot), Phospho-STAT3 (Tyr705, sc-8059, for immunoprecipitation), STAT3 (sc-482), IRF1 (CST, 8478), IRF3 (CST,11904), RELA (sc-372), MDA5 (CST, 5321), HCV NS3 (Abcam, ab13830), HCV Core (Thermo Fisher, MA1-080), HA (sigma, H9658), FLAG (WAKO, 014-22383), gp130 (sc-655), IL6Rα (sc-661), EGFR (Bioworld, BS3605), MERTK (sc-365499), FGFR1 (CST, 9740), IGF1Rβ (CST, 3018), β-actin (CST, 4970) and GAPDH (CST, 2118). S3I-201 (S1155) and Cryptotanshinone (S2285) were obtained from Selleck. Recombinant Human IL-29 (1598-IL-025), IFN-β (8499-IF-010) and IL-6 (206-IL-010) were obtained from R&D systems. IL-29/IL-28B (DY1598B) and IFN-beta (DY814) DuoSet ELISA kits as well as Ancillary Reagent Kit 2 (DY008) were purchased from R&D systems. Dual Luciferase Assay Kit (E1960) was purchased from Promega. Normal human liver sample was obtained from Liver Transplant Center, The Third Affiliated Hospital of Sun Yat-sen University.

### Cell lines

Huh7.5.1 was described^40^. HepG2 and Huh7 cell lines were obtained from Cell Bank of Chinese Academy of Sciences (Shanghai). 293FT cell line was obtained from Invitrogen. miR-122-Tet-On HepG2 cell line was generated using lentiviral vectors. Briefly, the transfer vector for expression of miR-122 hairpins (Supplementary Fig.6a) was generated by insertion of the TRE-3G promoter (*P*_TRE3G,_ from pTRE3G-IRES, Clonetech), GFP-miR-122x4 cassette^56^, PGK promoter (*P*_PGK_) and Neomycin resistance gene into the transfer vector of a third-generation lentiviral vector system. The transfer vector for expression of Tet3G (Supplementary Fig.6a) was generated by insertion of EF1α promoter (*P*_EF1A_), Tet3G (from pCMV-Tet3G, Clonetech), internal ribosome entry site (IRES) and Blasticidin resistance gene into another transfer vector. Lentiviruses were generated in 293FT cells according to the manual of ViraPower Lentiviral Packaging Mix (Invitrogen, K4975-00). HepG2 cells were infected with lentiviruses one after another, selected with G418 (1000 μg/ml) and Blasticidin (10 μg/ml) until cells do not die any more.

### Viral constructs and viral RNA preparation

pJFH1 and pSGR-JFH1 were described^42^. pJFH1-M was generated by the introduction of two point mutations (A-T) in the miR-122 binding sites on pJFH1. Viral RNAs were prepared by *in vitro* transcription (Ambion, AM1333). HCV RNAs were prepared as described^40^.

### Constructs

STAT3, NF-κB, IRF1 and ISRE reporter vectors were included in Cignal Finder™ Multi-Pathway Reporter Array (SABiosciences, CCA-901L). Eleven IRF1 promoter or enhance constructs were generated based on pGL3-basic (Promega). Mutations were introduced by PCR. Detailed information on the position of DNA fragment can be found in Supplementary Table 1. Reporter vectors for miR-122 binding sites were generated as described^56^. Briefly, 47-nt DNA fragments containing the putative binding site for miR-122 of each gene (5′-end 29-30 nt flanking sequences, 7–8 nt seed-matched sequences, 3′-end 10-nt flanking sequences) were cloned into Xho I/Not I-digested psiCHECK-2 vector (Promega) in forward direction. Mutated constructs were generated by replacing the miR-122 seed-matched sequence (CACTCC) with (GTGAGG). The coding sequences of protein genes were amplified by RT-PCR and cloned into pcDNA3 (IRF1), pRKW2-HA (HA-IRF3, HA-RELA, HA-NFKB1, HA-JUN and HA-ATF2), pRKW2-Flag (FL-EGFR, FL-MERTK, FL-FGFR1, FL-IGF1R, FL-ABL2 and FL-PKM2), or pcDNA3.1-HA (HA-IRF1, HA-JAK1, HA-JAK3, HA-IL1RN, HA-EPO, IGF2-HA and HA-FGF11). pcDNA3-RFP and pRKW2-HA-GFP were used as controls. All oligonucleotides used in this work can be found in Supplementary Table 5.

### Analysis of STAT3 binding sites on IRF1

ChIP-seq binding sites for STAT3 was analyzed on UCSC genome browser (GRCh37/hg19) using the “ENCODE Regulation Super-track”, and Txn Factor ChIP data (161 Transcription Factor) was selected. Seven STAT3 binding clusters within IRF1 locus (Supplementary Fig. 3a) were obtained and selected for ChIP-PCR assays (Supplementary Table 1). The STAT3 binding sites within these clusters were predicted by JASPAR (http://jaspar.genereg.net/). ChIP was performed using SimpleChIP^®^ Kit (CST, 9003) according to the manual. HepG2 cells from four 10-cm dishes were used in ChIP. Cells were starved overnight and treated with IL-6 (25 ng/ml) for 30 minutes before harvest. Totally 500μl cross-linked chromatin were obtained, and each 100μl (diluted into 500μl by adding 400μl 1X ChIP buffer) was immune-precipitated with 2 μg of the following antibodies: Normal Rabbit IgG (CST, 2729), Phospho-STAT3 (Tyr705, sc-8059), STAT3 (sc-482) and RELA (sc-372). After determination of the appropriate amplification cycle numbers by qPCR, the IPed chromatins were then analyzed by standard PCR with AccuPrime^TM^ PFX SuperMix (Invitrogen, 12344-040).

### Collection of Candidate STAT3 Activators

We totally collected 330 candidates (known and potential) that can be detected in our microarray data (Supplementary Table 3), by strategies as follows:

1. In the website of R&D systems (https://www.rndsystems.com/cn/pathways/jak-stat-signalling-pathway), there is a comprehensive collection on the cytokines & receptors that have been involved in JAK-STAT signalling pathway, which includes: JAKs, IL6 family & receptors, IL10 family & receptors, IL12 family & receptors, interferons & receptors, receptor tyrosine kinases (RTKs) & ligands, G Protein-Coupled Receptors (GPCRs) & ligands, Homodimeric Hormone Receptors (HHRs) & ligands, Common beta Chain Receptor Family (CbCRF) & receptors, Common gamma Chain Receptor Family (CgCRF) & receptors. All these genes were collected.
2. Komurov and colleagues have performed high-throughput siRNA-mediated screening on genes supporting EGF-dependent signalling [Supp. Ref. 1] and identified many genes could regulate STAT3 tyrosine phosphorylation. All the genes promoting STAT3 phosphorylation (Other Known Regulator) were collected.
3. Many RTKs and non-receptor tyrosine kinases (N-RTKs) could regulate STAT3 phosphorylation [Supp. Ref. 2-9]. Therefore, we collected all known RTKs (20 families) [Supp. Ref. 10] as well as known N-RTKs (according the annotation in GeneCards).
4. Given that interleukins are involved in diverse immune activities, we arbitrarily collected all other interleukins (Other Cytokine) and their receptors (Other Cytokine Receptor), as well as related binding proteins (Cytokine-receptor interaction) as potential STAT3 regulators.
5. We also collected potential RTKs, ligands, as well as RTK-related genes based on their names.

### Cell cultures and transfections

HepG2 and miR-122-Tet-On HepG2 cells were maintained in MEM supplemented with 10% fetal bovine serum (FBS) and antibiotics (100 U/ml penicillin and 100 μg/ml streptomycin). PC3 was maintained in F-12K (Sigma, N3520) supplemented with 10% FBS and antibiotics. Huh7-derived cell lines were maintained in DMEM supplemented with 10% FBS and antibiotics.

Mimics and siRNAs were reversely transfected into cells using Lipofectamine^TM^2000 (Invitrogen) at a final concentration of 20nM unless otherwise indicated (for more than one siRNAs, the concentration represents all mixed siRNA oligos). HCV RNAs were transfected into HepG2 cells using TransIT^®^-mRNA Transfection Kit (Mirus), into Huh7 and Huh7.5.1 cells using Lipofectamine^TM^2000, at a final concentration of 1 μg/ml. Low-molecular-weight Poly(I:C) (Invivogen) were transfected into HepG2 or Huh7 cells using Lipofectamine^TM^2000 at a final concentration of 2 μg/ml. For PC3 cells, we transfected 0.4μg/ml Poly(I:C) because high amount of Poly(I:C) could induce serious cell death. Plasmids were transfected into HepG2, Huh7 and PC3 using ViaFect™ Transfection Reagent (Promega), into 293FT using Lipofectamine^TM^2000.

For reporter assays on signalling pathways, HepG2 cells were transfected in 48-well-plates, with 20 nM mimics, 0.1 μg firefly luciferase reporter plasmids (transcription factor responsive), and 0.0025 μg control plasmids (constitutively expressing Renilla luciferase); 48 hours later, cells were further transfected with SGR-JFH1 RNA (0.25 μg) for the indicated treatment durations. For comparison of 11 different IRF1 promoter/enhancer constructs, HepG2 cells were transfected in 48-well-plates, with 0.2 μg Firefly luciferase reporter plasmids and 0.005 μg control Renilla plasmids; 48 hours later, cells were further transfected with poly(I:C) for 24 hours. For assessment of the effects of STAT3 and other transcription factors on IRF1 constructs, 293FT cells were transfected in 48-well-plates, with 0.1 μg reporter plasmids, 0.1 μg transcription factor expressing plasmids, and 0.0025 μg control Renilla plasmids. For miR-122 target identification, 293FT cells were transfected in 96-well-plates, with 0.05 μg reporter constructs, 0.05 μg miR-122 or miR-neg expressing vectors, as previously described^56^.

### qRT-PCR

Total RNA was isolated with Trizol reagent (Invitrogen). The expression of mRNAs, HCV RNA and miR-122 were quantified by SYBR Green-based Real-time PCR. Reverse transcription (RT) reactions for mRNAs were done using GoScript™ RT System (Promega, A5000), with random primers and oligo dT. RT reactions for miR-122 were done using PrimeScript™ RT reagent Kit with gDNA Eraser (TaKaRa, 047A), with a stem-loop primer. RT reactions for HCV RNA were performed using SuperScript^®^ III Reverse Transcriptase, with a specific primer. qPCRs were performed with GoTaq^®^ qPCR Master Mix (Promega, A6002).

### Western blots

Total protein of cell lines and liver tissues were prepared using RIPA buffer (50mM Tris-HCl, pH 8.0, 150mM Sodium chloride, 1% NP-40, 0.5% sodium deoxycholate, 0.1% sodium dodecyl sulfate (SDS), 2mM EDTA) with 1X Protease Inhibitor Cocktail (Roche, 04693132001) and 1x PhosSTOP™ phosphatase inhibitor (Roche, 04906837001). Equivalent total protein extracts were separated by SDS-polyacrylamide gel electrophoresis (PAGE) and transferred onto Protran™ nitrocellulose (NC) membranes (Amersham, 10600001). The blots were incubated first with a primary antibody and then, with horseradish peroxidase-conjugated secondary antibody (CST, 7074 or 7076). Immunoreactive bands were visualized by using Luminata Crescendo Western HRP substrate (Millipore, WBLUR0500) and exposed to X-ray films.

### Microarray

HepG2 cells were transfected with 50 nM control or miR-122 mimics and cells were harvested 72 hours after transfection. 1000ng/ml doxycycline was added into the culture medium of miR-122-Tet-On HepG2 cells for 72 hours to induce miR-122 expression; Cells without doxycycline served as control. RNAs were sent to microarray analysis with GeneChip^®^ Human Genome U133 Plus 2.0 Arrays (CapitalBio Corporation). Microarray data are deposited in GEO with accession number GSE99663. In order to get enough genes for our study, we took a 20% threshold (up- or down-regulation) because the change in most of candidate STAT3 activators was not very apparent. e.g., setting a 50% cut-off (generally used in analyzing microarray data) resulted in zero gene left. Besides, as we noticed that the different probe sets for the same gene sometimes gave dissimilar results, we selected candidate genes for further qRT-PCR validation as long as there was a significant difference in the signals from any probe for a given gene.

### Statistical Analysis

Data are representative of at least three independent experiments. For statistical analysis, the differences were analyzed using two-sided Student’s *t*-tests when comparing the effect of two factors on the same subjects, or one-way analysis of variance (ANOVA) when comparing the effect of multiple factors. A *P* value < 0.05 was considered statistically significant. \**P* < 0.05, \*\**P* < 0.01 and \*\*\**P* < 0.001. Error bars represented standard error of the mean (SEM) in qRT-PCR and ELISA data, or standard deviation (SD) in luciferase data.

## ACKNOWLEDGMENTS

We thank Takaji Wakita (National Institute of Infectious Diseases, Japan) for use of pJFH1 and pSGR-JFH1, Francis Chisari (Scripps Research Institute, US) for use of Huh7.5.1 cell line, Jin Zhong (Institut Pasteur of Shanghai, China) for providing these materials. We thank Hong-bing Shu (Wuhan University, China) for providing the FLAG and HA-tagged RKW2 vectors. We thank Xiang-Dong Fu (University of California, San Diego) and Jun Cui (Sun Yat-sen University, China) for helpful discussions. This research was supported by the National Natural Science Foundation of China (No. 31200593, 31230042 and 31471223), the Natural Science Foundation of Guangdong Province (2014A030313163), the project of Science and Technology of Guangzhou (201504010022) and the National Basic Research Program of China (2011CB811300).

### AUTHOR CONTRIBUTIONS

H.X. and S. Xu conceived and designed the experiments; S. Xu, H.X., S. Xie, Y.Z., W.Z., and M.Z. performed experiments; H.X. and S. Xu analyzed data; J.Y. assisted in miR-122 target prediction and microarray data analysis; L.Q. and H.Z. supervised the study. H.X., S. Xu and L.Q. wrote the manuscript. All authors edited the manuscript.

### COMPETING FINANCIAL INTERESTS

The authors declare no competing financial interests.

